# Opposing transcriptional mechanisms regulate *Toxoplasma* development

**DOI:** 10.1101/094847

**Authors:** Dong-Pyo Hong, Joshua B. Radke, Michael W. White

## Abstract

The *Toxoplasma* biology that underlies human chronic infection is developmental conversion of the acute tachyzoite stage into the latent bradyzoite stage. We investigated the role of two alkaline-stress induced ApiAP2 transcription factors, AP2IV-3 and AP2IX-9, in bradyzoite development. These factors were expressed in two overlapping waves during bradyzoite development with AP2IX-9 increasing expression earlier than AP2IV-3, which peaked as AP2IX-9 expression was declining. Disruption of the AP2IX-9 gene enhanced, while deletion of AP2IV-3 gene decreased tissue cyst formation demonstrating these factors have opposite functions in bradyzoite development. Conversely, conditional overexpression of FKBP-modified of AP2IX-9 or AP2IV-3 with the small molecule Shield 1 had a reciprocal effect on tissue cyst formation confirming the conclusions of the knockout experiments. The AP2IX-9 repressor and AP2IV-3 activator tissue cyst phenotypes were borne out in gene expression studies that determined many of the same bradyzoite genes were regulated in an opposite manner by these transcription factors. A common gene target was the canonical bradyzoite marker, BAG1, and mechanistic experiments determined that like AP2IX-9, AP2IV-3 regulates a BAG1 promoter-luciferase reporter and specific binds the BAG1 promoter in parasite chromatin. Altogether, these results suggest the AP2IX-9 transcriptional repressor and AP2IV-3 transcriptional activator likely compete to control bradyzoite gene expression, which may permit *Toxoplasma* to better adapt to different tissue environments and select a suitable host cell for long term survival of the dormant tissue cyst.

**IMPORTANCE:** *Toxoplasma* infections are life-long due to the development of the bradyzoite tissue cyst, which is effectively invisible to the immune system. Despite the important clinical consequences of this developmental pathway, the molecular basis of the switch mechanisms that control formation of the tissue cyst is still poorly understood. Significant changes in gene expression are associated with tissue cyst development and ApiAP2 transcription factors are an important mechanism regulating this developmental transcriptome. However, the molecular composition of these ApiAP2 mechanisms are not well defined and the operating principles of ApiAP2 mechanisms are poorly understood. Here we establish that competing ApiAP2 transcriptional mechanisms operate to regulate this clinically important developmental pathway.

## INTRODUCTION

The molecular switch mechanisms that control bradyzoite development have long been sought, but have remained elusive as have clinical treatments effective against the tissue cyst stage. The life cycle of *Toxoplasma* is heteroxenous with a sexual reproductive cycle exclusive in the felid host and a second clonal reproductive phase (intermediate life cycle) that is possible in virtually any endothermic animal. There are two competing demands of the *Toxoplasma* intermediate life cycle; expand tachyzoite numbers to ensure survival and production of the bradyzoite-tissue cyst required to transmit the infection to the next host. Our previous studies demonstrate *Toxoplasma* bradyzoite differentiation is a multi-step process requiring slowing of the tachyzoite cell cycle combined with key changes in gene expression and finally exit from the cell cycle (1). Epigenetic and gene-specific transcription mechanisms are implicated (2) indicating a transcriptional network regulates these developmental processes. However, the molecular basis for these controls is poorly defined and the *Toxoplasma* factors required are not completely identified.

Based on *“in silico”* analysis, Balaji *et al.,* (2005) proposed there is a major lineage of transcription factors in Apicomplexa (ApiAP2 factors) that are weakly similar to a group of transcription factors found in plants (3). The plant-like transcription factors in the Apicomplexa harbor one or up to six Apetala2 (AP2) DNA binding domains (4). The substantial majority of ApiAP2 domains have a single characteristic subtype of the AP2 structural fold. This is determined by a sequence pattern in the DNA binding domain (~60 amino acids) that is strongly similar in the different apicomplexan lineages and has distinct differences from the plant AP2 patterns (5).There is now solid evidence ApiAP2 factors are bona vide transcription factors regulating parasite gene expression through direct binding of specific promoter elements and many Apicomplexa genes transcribed in specific life cycle stages are regulated by this type of mechanism (6-9). For some ApiAP2 factors, the influence on gene expression may be indirect through binding and modifying the activity of other chromatin factors (10). There are 67 total ApiAP2 genes encoded in the *Toxoplasma* genome with product names assigned by chromosome location and order (e.g. AP2IV-3 searchable at ToxoDB)(11–13). In other Apicomplexa lineages there are far fewer ApiAP2 genes (e.g. 27 in *Plasmodium falciparum* (3), 17 in *Cryptosporidium spp,* (14)), which may reflect the unique evolution of this protozoan family of parasites (15). Recent comparative genome analysis of lineage-specific apicomplexan species with the free-living relatives *Chromera velia* and *Virella brassicaformis* reveals a large expansion of AP2-like factors likely occurred prior to the splitting of the Chromerids and Apicomplexa (15). Thereafter, the ApiAP2 gene family diverged independently in each descending Apicomplexa lineage. For example, it is very likely that the smaller number of ApiAP2 factors present in modern *Plasmodium spp* parasites is the result of at least three major reductions in gene content since the Apicomplexa split from the Chromerids (15). These gene reductions may have had other functional consequences for the evolution of ApiAP2 mechanisms as so called master regulators of parasite development equivalent to AP2-G in *P. falciparum* (16) have yet to be identified in *Toxoplasma* or other coccidians where regulation of developmental gene expression is potentially more complex. Our studies here expand our understanding of how ApiAP2 factors regulate development of the clinically important *Toxoplasma* tissue cyst and establish there are multiple opposing transcriptional forces competing to preserve asexual replication or shift development towards the formation of the transmissible cyst stage. These ApiAP2 repressors and activators are responsive to stress-signals and are active early in tachyzoite to bradyzoite differentiation where they regulate many of the same parasite genes. Our results demonstrate that transcription factors regulating tachyzoite to bradyzoite development are acting at the level of the individual parasite and are not coordinated by the intravacuole environment, which may help explain the stochastic nature of this developmental pathway.

## RESULTS

### AP2IV-3 is one of five ApiAP2 factors expressed early in bradyzoite development

New transcriptome datasets generated from stages across the full *Toxoplasma* life cycle including the feline sexual cycle has permitted the reanalysis of ApiAP2 factors (Tables 1 and S1). We selected datasets deposited in ToxoDB (http://toxodb.org/toxo/) representing Type II strain transcriptomes from *in vitro* tachyzoites, early and mature bradyzoites, merozoites, and sporozoites to assess how all 67 ApiAP2 mRNAs are developmentally expressed (ranked high to low, 1-67 in each sample). For 22 ApiAP2 genes the mRNA expression data did not identify a specific developmental profile. Many non-developmental ApiAP2 factors are expressed at low levels (<50th percentile, 18 TL), while others showed variable moderate to higher expression across the stages. Notably, the cell cycle regulated tachyzoite factor, AP2VI-1 (12), was the only ApiAP2 mRNA expressed at high levels (ranked 6th or higher) across all the transcriptomes examined (Table S1). Relatively few ApiAP2 genes were exclusively or highly expressed in a single life cycle stage, whereas multiple life cycle stage expression was more common (Table 1). The expression of ten ApiAP2 mRNAs were distinctly higher in intermediate life cycles stages, however, many more ApiAP2s were expressed in these stages plus oocyst environmental stages (10 TL) or feline cycle stages (12 TL)(Table 1). Similar ApiAP2 expression profiles in tachyzoites and bradyzoites was expect because the transcriptomes of these stages are highly correlated (17, 18). Common ApiAP2 expression between the intermediate life cycle and oocyst stages was also anticipated as more than 20 years ago we established the principal that coccidian sporozoites are pre-programmed for the next stage of development (19), which in *Toxoplasma* is the tachyzoite. The expression of ApiAP2 factors during bradyzoite development revealed surprising patterns. A total of six ApiAP2 mRNAs (AP2Ib-1, AP2IV-3, AP2VI-3, AP2VIIa-1, AP2VIII-4, and AP2IX-9) were strongly up regulated in low passage Type II ME49 parasites by 2 days following alkaline-stress induction of tachyzoites and four of these factors were down regulated in mature bradyzoites (21 day murine brain tissue cysts) indicating their role is restricted to the transition of the tachyzoite to the bradyzoite. The expression of three of these ApiAP2 factors during early bradyzoite development has been confirmed at the protein level (AP2IV-3, Fig. 1A; AP2Ib-1, not shown; AP2IX-9, ref. 6). In this analysis, no ApiAP2 was determined to be exclusively up regulated in mature bradyzoites, where instead the dominant phenotype was the down regulation of eleven ApiAP2 mRNAs (Table 1) expressed at higher levels in tachyzoites and/or early bradyzoites. The samples analyzed here are population snap shots and may miss more transiently regulated ApiAP2 factors such as those that are cell cycle regulated (12) and also sexual stage ApiAP2s as these stages were not specifically sampled. In addition, there are strain-specific differences known to affect ApiAP2 expression in *Toxoplasma* (20) that were evident but not further addressed here.

**FIG 1.**
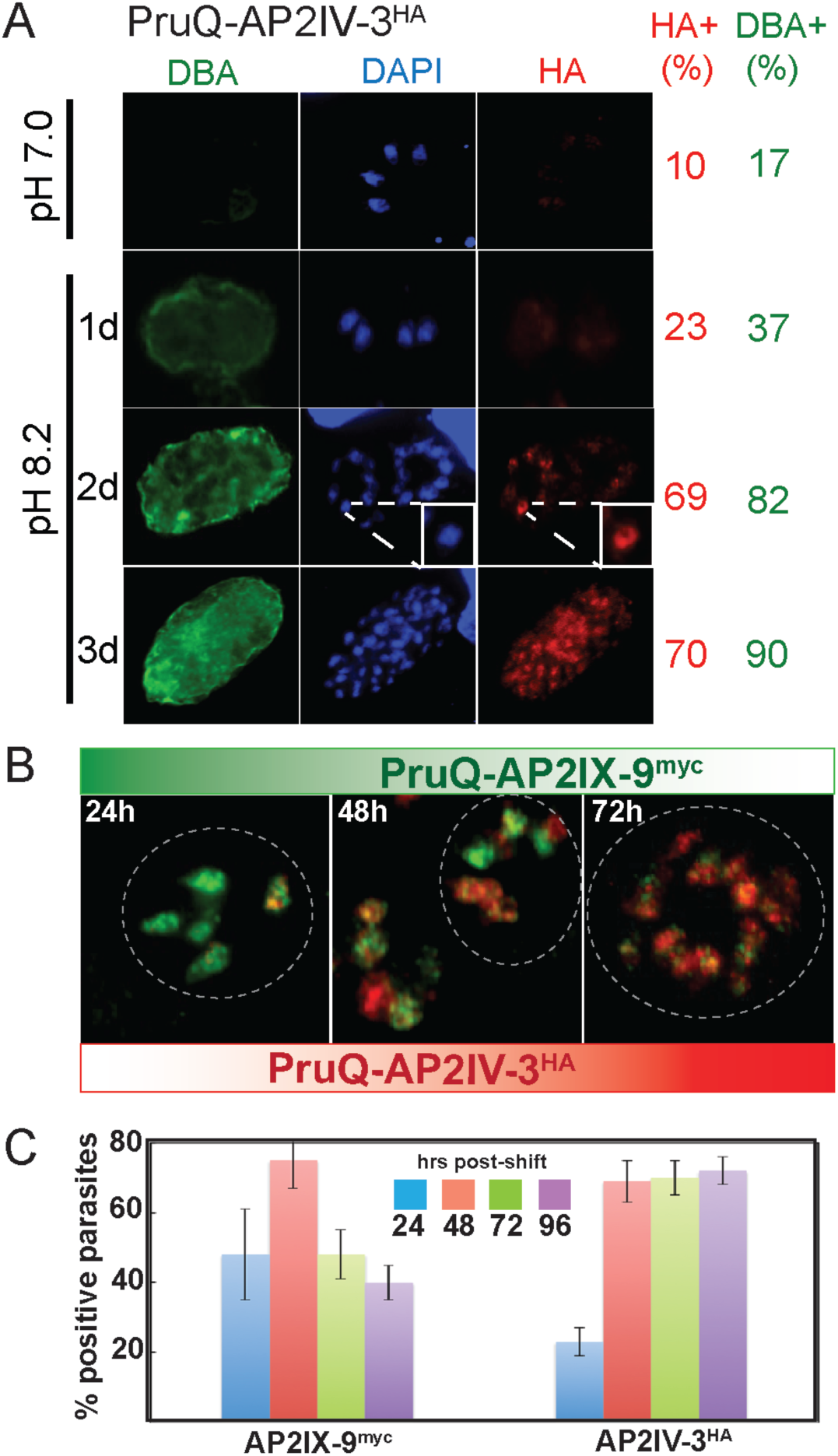
*Toxoplasma* AP2IV-3 is induced by stress during bradyzoite development. (A) The ApiAP2 factor, AP2IV-3, was C-terminally epitope tagged by genetic knock-in at the endogenous locus with 3xHA in the PruQ strain. PruQ-AP2IV-3^HA^ parasites were propagated in human foreskin fibroblasts (HFF) on cover slips using pH 7.0 or pH 8.2 media. A various times cover slips were fixed and costained with anti-HA antibody (red, for AP2IV-3 expression), biotin-labeled *D. biflorus* agglutinin (DBA, green for the presence of cyst wall), and DAPI (blue for genomic DNA). HA- or DBA-positive parasites (indicated on the right) were quantified in 100 randomly vacuoles. The inset Dapi and anti-HA images of parasites at day 2 post-pH 8.2 media shift show co-localization indicating AP2IV-3^HA^ was exclusively nuclear. (B and C) Utilizing sequential genetic knock-in, AP2IX-9 (3xmyc) and AP2IV-3 (3xHA) were C-terminally epitope tagged at the endogenous loci in the PruQ parent strain. IFA analysis of the resulting dual-tagged transgenic clone was performed using anti-HA (red stain) and anti-myc (green stain) monoclonal antibodies. Representative vacuoles at 24, 48 and 72 post-alkaline shift (pH 8.2 media) are shown. The dashed circles indicate vacuole boundaries. Triplicate quantitative analysis of AP2IX-9^myc^ and AP2IV-3^HA^ positive parasites was determined in 100 randomly selected vacuoles over 96 hours post-shift into pH 8.2 media.

**Table 1.** Categories of developmental ApiAP2 mRNA expression.

| Specific Developmental Expression | *Toxoplasma* ApiAP2 Name |
| --- | --- |
| **intermediate cycle**<br>(at least one stage) | AP2V-1, AP2VIIa-2, AP2VIIa-5, AP2VIIb-3<sup>1</sup>, AP2VIII-4, AP2IX-4<sup>1</sup>, AP2IX-9, AP2XI-2<sup>1</sup>, AP2XI-4<sup>1</sup>, AP2XII-8 |
| **feline cycle**<sup>2</sup> | AP2III-4, AP2IX-6 |
| **environmental stages**<sup>3</sup> | AP2III-3, AP2VI-2, AP2VIIa-7, AP2X-2, AP2XII-3, |
| **feline cycle and environmental stages** | AP2IV-1, AP2IX-1, AP2X-10 |
| **intermediate cycle plus sporozoites** | AP2III-2, AP2IV-5, AP2VIIa-1, AP2VIIa-2, AP2VIIa-3, AP2VIIa-6, AP2VIIa-8, AP2VIII-3, AP2X-5, AP2X-7 |
| **intermediate cycle plus merozoites** | AP2Ib-1<sup>4</sup>, AP2VI-3<sup>4</sup>, AP2VIIa-4, AP2VIIa-9, AP2VIII-5, AP2VIII-6, AP2VIII-7, AP2IX-6, AP2IX-8, AP2X-4, AP2X-9, AP2XII-6 |
| **up regulated in early bradyzoites** | AP2Ib-1<sup>5</sup>, AP2IV-3<sup>5</sup>, AP2VI-3, AP2VIIa-1, AP2VIII-4, AP2IX-9<sup>5</sup> |
| **down regulated in mature *in vivo* bradyzoites,** | AP2Ib-1<sup>4</sup>, AP2IV-3<sup>4</sup>, AP2IV-4, AP2IV-5, AP2V-1, AP2VI-3<sup>4</sup>, AP2VIIa-5, AP2VIIa-6, AP2VIII-5, AP2IX-9<sup>4</sup>, AP2XII-6 |
| **Comments** | <sup>1</sup>all intermediate life cycle stages<br><sup>2</sup>day 3, 5, 7 post-infection of the cat host, see ToxoDB<br><sup>3</sup>unsporulated oocysts (zygote) to fully sporulated oocysts containing sporozoites<br><sup>4</sup>also specific for early bradyzoites<br><sup>5</sup>confirmed at the protein level (see Table S1) |

AP2IV-3 is one of the six ApiAP2 factors up regulated in early bradyzoites (Table 1) with AP2IV-3 mRNA significantly increased in comparison to tachyzoite expression in response to 48 h alkaline-stress in Type II (71st percentile), and Type III (84th) strains (see AP2IV-3 gene record, ToxoDB). AP2IV-3 mRNA expression in 21 day bradyzoites from mouse brain (18) is substantially down regulated (Table S1). Epitope tagging of AP2IV-3 in a Prugnauid genetic background (Pru-*Δku80Δhxgprt,* designated as PruQ) by genetic knock-in revealed a nuclear factor that was induced by alkaline-stress during the early stages of bradyzoite switching, while largely absent from tachyzoites (Fig. 1A).

In order to better understand AP2IV-3 protein expression, we generated a PruQ transgenic strain by sequential knock-in that expresses dual epitope-tagged ApiAP2 factors, AP2IV-3^HA^ and AP2IX-9^myc^ (Fig. 1B and C). AP2IX-9 is also one the five early expressed bradyzoite ApiAP2 factors that is down regulated at the mRNA and protein levels in mature bradyzoites (6). Immunofluorescence (IFA) analysis of the dual tagged strain revealed that AP2IX-9^myc^ expression increased before AP2IV-3 following the change to pH 8.2 media, while AP2IV-3^HA^ expression became dominant later in development as AP2IX-9^myc^ expression declined (Fig. 1C). AP2IX-9^myc^ and AP2IV-3^HA^ expression patterns were neither synchronous nor exclusive and parasites sharing a vacuole could be dual or single expressers of these factors (Fig. 1B). The heterologous expression profiles of AP2IX-9 and AP2IV-3 were similar to expression patterns of other bradyzoite proteins (e.g. BAG1, LDH2, SRS9) that reflect the characteristic asynchronous nature of bradyzoite development (21).

### Genetic disruption and overexpression of AP2IV-3 or AP2IX-9 demonstrate opposite functional roles in regulating tissue cyst formation

In order to understand the function of AP2IV-3, we isolated AP2IV-3 knockout clones produced in the Pru strain (Pru=Prugnauid-*Δhxgprt*) and determined how well Pru-*Δap2IV-3* parasites were able to form tissue cysts in alkaline media. We compared these results to the Pru parent and also to a Pru transgenic clone in which we disrupted the AP2IX-9 gene (Pru-*Δap2IX-9*)(Fig. 2B). The Pru parent spontaneously forms 15-20% tissue cysts in standard pH 7.0 media (Fig. 1A) similar to the pH 7.4 media (Fig. 2B, blue bars). The deletion of the AP2IX-9 gene in the Pru strain increased tissue cyst formation to levels higher than the Pru parent under mild alkaline-stress conditions (Fig 2B, blue and red bars, pH 7.4 and pH 7.8 media). These results are consistent with our previous report that AP2IX-9 is a bradyzoite transcriptional repressor (6). By contrast, deletion of the AP2IV-3 gene resulted in a lower capacity to form tissue cysts in pH 7.4 and pH 7.8 media (Fig. 2B). Thus, culturing both transgenic parasites in pH 7.8 media induced Pru-*Δap2IX-9* parasites to form tissue cysts (68%) at more than twice the rate as the Pru-*Δap2IV-3* strain in this media (29%). Complementation of the Pru-*Δap2IV-3* strain with a cosmid carrying a single copy of the AP2IV-3 gene fully restored the tissue cyst formation to levels that matched or exceeded the Pru parent or Pru-*Δap2IX-9* parasites (Pru-*Δap2IV-3*::AP2IV-3 cyst numbers, pH 7.4=43%, pH 7.8=72%, pH 8.2=96% compare to Fig. 2B). A reduction of tissue cyst capacity was also observed when the AP2IV-3 gene was disrupted in a Type III CTG genetic background (Fig. S1, pH 8.2 media for 72 h). The reduction in tissue cyst formation caused by the loss of AP2IV-3 was more pronounced in CTG than in the Pru strain, which shows higher rates of spontaneous differentiation than CTG and is more readily induced to develop (compare blue bars in Figs. 2B and 3). Notwithstanding the strain differences, the results with Pru and CTG transgenic strains demonstrate AP2IV-3 functions in a manner opposite to AP2IX-9 (6) in promoting tissue cyst formation.

**FIG 2.**
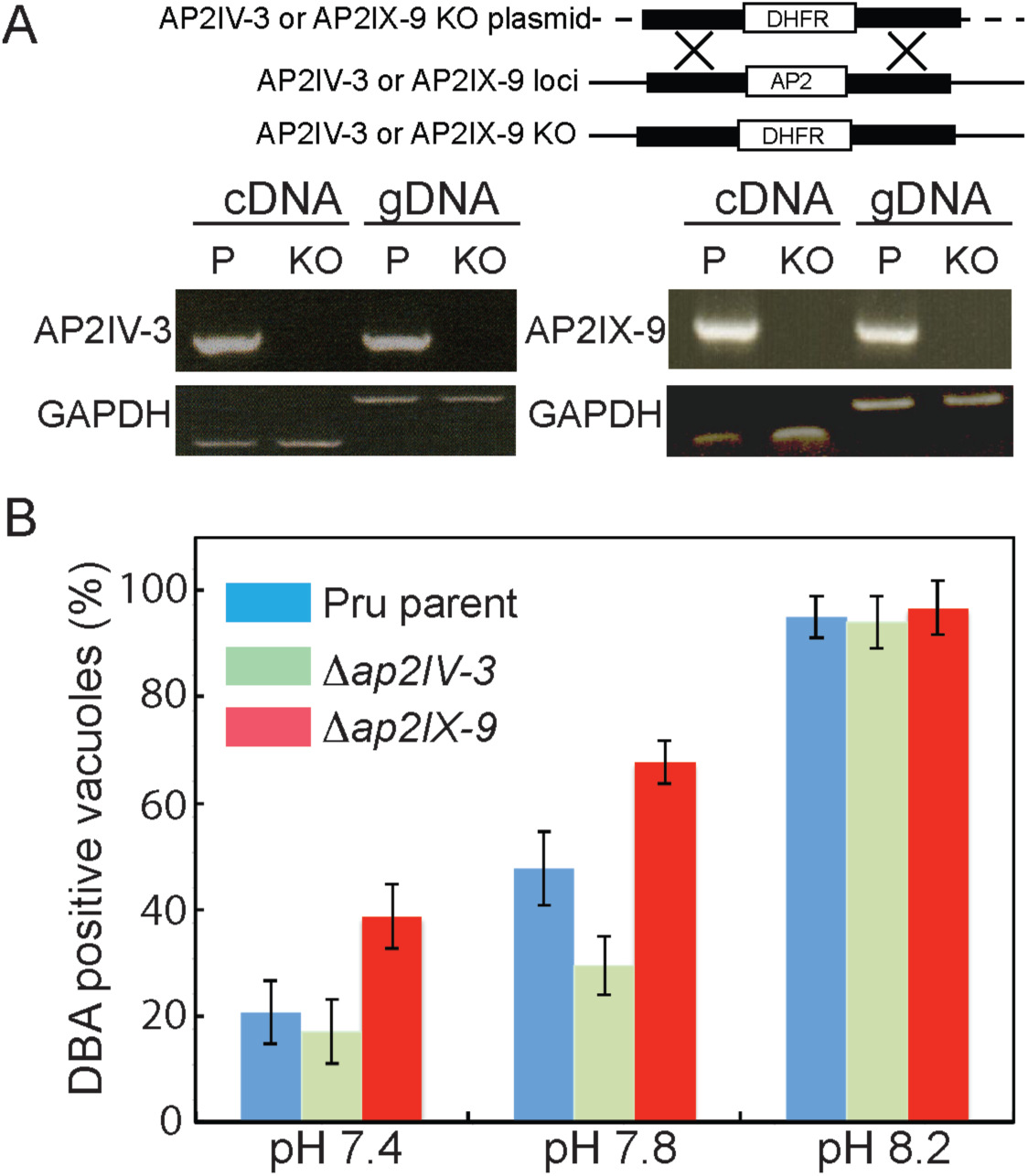
Disruption of AP2IV-3 and AP2IX-9 genes alters the capacity to form tissue cysts. (A) Schematic representation of knockout constructs for the AP2IV-3 or AP2IX-9 locus. Utilizing the pyrimethamine resistant DHFR selectable marker and CRISPR methodology disruptions of AP2IV-3 and AP2IX-9 were achieved as shown by the PCR analysis of transgenic DNA (gDNA) and RT-PCR of complementary DNA (cDNA) for parent (P) and knockout (KO) strains; proper fragment sizes cDNA (202 bp) and gDNA (634 bp). PCR analysis of GAPDH was included as a negative control. (B) Tissue cyst formation of the Pru strain parent versus Pru-*Δap2IV-3* or Pru-*Δap2IX-9* knockout clones grown under different alkaline-stress conditions. The proportion of DBA-positive vacuoles in 100 randomly selected vacuoles was assayed in triplicate. Note that disruption of AP2IX-9 (red bars) leads to greater tissue cyst formation in milder alkaline conditions, while the opposite result was observed when the AP2IV-3 gene was deleted (green bars).

**FIG 3.**
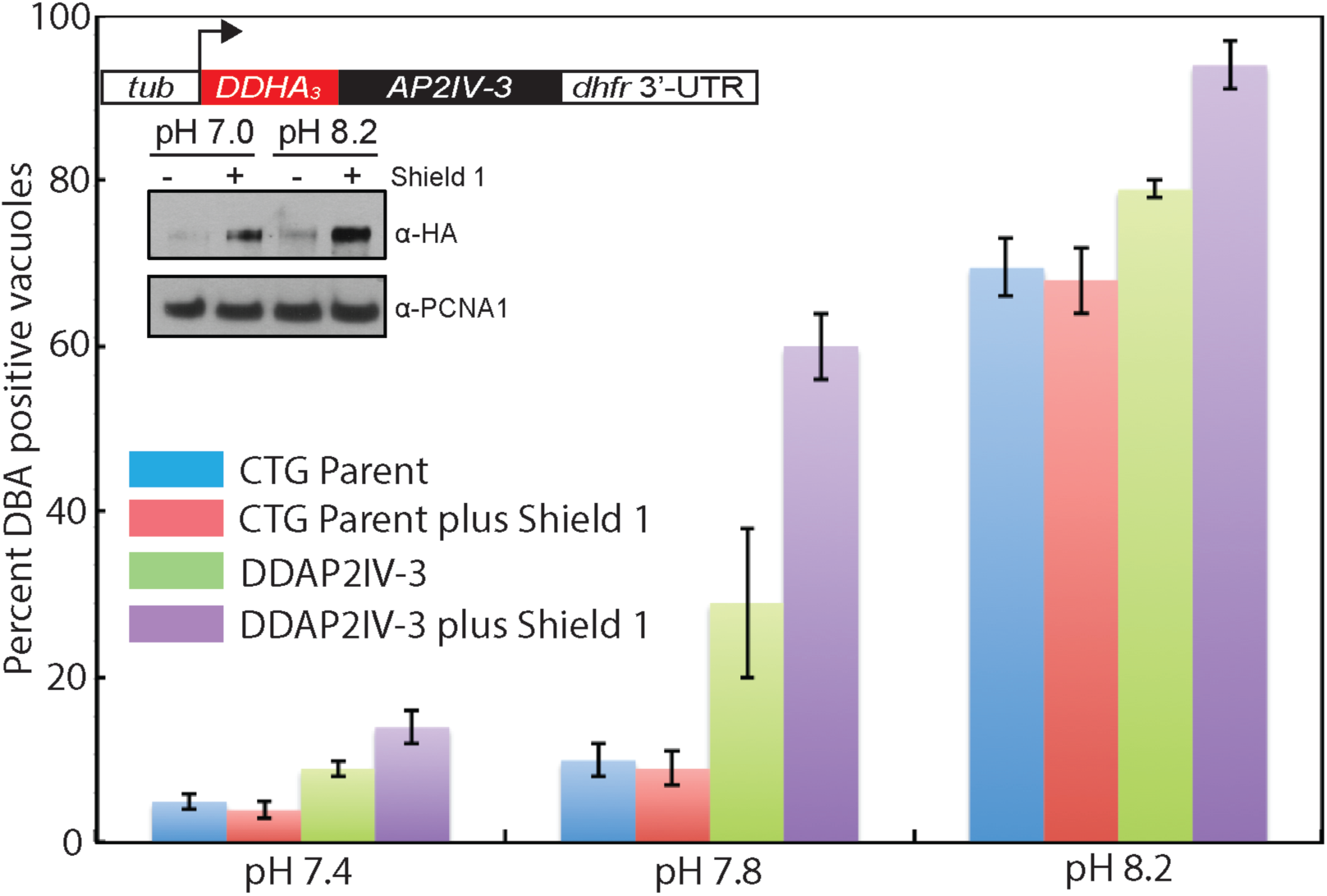
Overexpression of AP2IV-3 enhances tissue cyst formation. The 72 h capacity to form tissue cysts of the Type III CTG parent strain compared to a CTG transgenic clone expressing ^DD^AP2IV-3 was evaluated after shift into mild to strong alkaline-media plus or minus Shield 1 (shld-1=100nM) as indicated. The fraction of DBA-positive parasites in 100 randomly selected vacuoles was measured in triplicate. Note that Type III CTG parental tachyzoites have a lower propensity to form tissue cysts spontaneously and are more resistant to milder alkaline pH media conditions than Pru parental strain tachyzoites (Fig 2B, blue bars).(Inset) Diagram of the ^DD^AP2IV-3 overexpression construct and Western blot of ^DD^AP2IV-3 expression with or without Shield 1. Note that ^DD^AP2IV-3 was induced to detectable levels by pH 8.2 media alone and to higher levels when Shield 1 was added to transgenic parasites grown in pH 7.0 and pH 8.2 media. The levels of nuclear TgPCNA1 as revealed by rabbit polyclonal antibody staining was included as a loading control.

To confirm the role of AP2IV-3 in promoting tissue cyst formation, we constructed an AP2IV-3 overexpression strain using the FKBP (DD)/Shield 1 conditional expression model (Fig. 3)(22–24). The FKBP peptide combined with three copies of the HA epitope tag was fused to the N-terminus of the AP2IV-3 coding region (Fig. 3, inset diagram and Western blot) and then transfected into the Type III CTG strain, which has a lower rate of spontaneous differentiation (Fig. 3, blue bars). A clonal CTG-^DD^AP2IV-3 isolate was exposed to increasing alkaline-stress plus or minus Shield 1 (100nM) and the tissue cyst number determined after 72 h (Fig. 3). As expected, overexpression of AP2IV-3 had the opposite effect on tissue cyst formation compared to the loss of AP2IV-3 by genetic knockout. Under mild alkaline media conditions (pH 7.4 and 7.8 media) tissue cyst formation of the CTG-^DD^AP2IV-3 strain was substantially increased in a Shield 1-dependent manner (Fig. 3). Combining stronger stress conditions (pH 8.2) with Shield 1 led to nearly 100% of CTG-^DD^AP2IV-3 parasites contained within DBA+-tissue cysts similar to the images shown in Figure 1A. This is one of the highest tissue cyst conversion rates that we have observed for any strain *in vitro.* By contrast, previous experiments where AP2IX-9 was overexpressed clearly demonstrated a strong repressive effect on tissue cyst formation in Type II and III strains (6).

### AP2IV-3 is an activator of bradyzoite gene expression

To identify the genes potentially controlled by AP2IV-3, duplicate RNA samples were prepared from CTG parent and CTG-^DD^AP2IV-3 and *CTG-Δap2IV-3* transgenic strains grown under different conditions of alkaline-stress plus 100nM Shield 1 (Fig. 4). Changes in mRNA expression due to Shield 1-induced overexpression of ^DD^AP2IV-3 parasites grown in pH 7.0 media revealed 42 genes (of 320 alkaline-responsive genes)(25) were affected when compared to CTG parental tachyzoites (compare Fig. 4A & B). A total of 36 genes were up regulated more than 1.5-fold, while only 6 genes were down regulated (see Dataset S1). Many of the genes up regulated by AP2IV-3 overexpression in tachyzoites are known bradyzoite genes that are induced by strong alkaline-stress in parental strains including the canonical bradyzoite markers BAG1 (in ^DD^AP2IV-3 tachyzoites up ~12 fold) and LDH2 (up ~4 fold) as well as a bradyzoite-specific rhoptry protein (26), BRP1 (up ~14 fold), and the SAG-related surface antigen SRS22A (up ~4 fold). A single gene encoding an unknown protein containing several ankyrin-repeats was the only strongly down regulated mRNA. Applying mild alkaline-stress conditions plus Shield 1 (pH 7.8 media) did not significantly alter the 42 putative AP2IV-3 target genes over Shield 1 stabilization of ^DD^AP2IV-3 in pH 7.0 media (Fig. 4C). By contrast, disruption of the AP2IV-3 gene largely led to the opposite affect on gene expression (Fig. 4D and Dataset S1). The majority of the 36 genes (24 of 36) increased by overexpression of ^DD^AP2IV-3 in tachyzoites failed to be induced in CTG-*Δap2IV-3* parasites shifted into pH 8.2 media, while these genes were induced in the CTG parent strain (Dataset S1). For example, BAG1 mRNA expression in alkaline-stressed CTG-*Δap2IV-3* parasites reached only ~10% the level of BAG1 mRNA induced by alkaline-stress of CTG parental parasites (Dataset S1). Intriguingly, a number of the genes regulated by AP2IV-3 are also controlled by AP2IX-9 (6)s including BAG1 and LDH2, although with opposite effects; overexpression of AP2IV-3 increased whereas elevated AP2IX-9 repressed mRNA expression (Fig. 4D). The mRNA encoding AP2IV-3 is also up regulated in merozoites isolated from cats and mature sporozoites (Table S1) suggesting this factor may have multiple roles in regulating gene expression across the *Toxoplasma* life cycle (17). Consistent with this idea, some of the genes up regulated by AP2IV-3 overexpression are also increased in merozoites and sporozoites (Dataset S2). For example, the rhoptry protein BRP1 (26) is elevated 14-fold by AP2IV-3 overexpression in tachyzoites and is ~10 fold higher in merozoites compared to tachyzoites of the same strain (Dataset S2).

**FIG 4.**
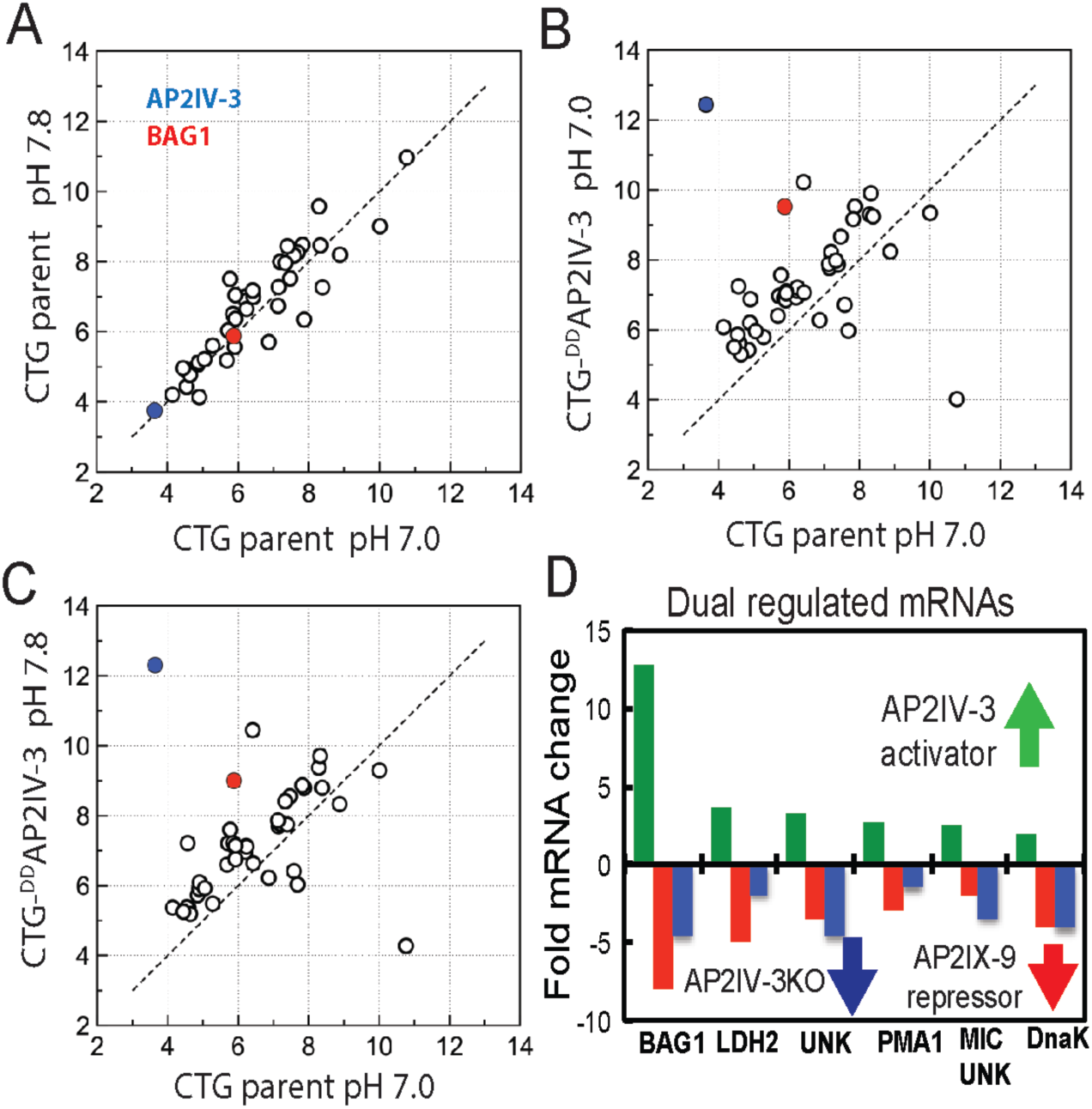
AP2IV-3 is an activator of bradyzoite gene expression. A total of 42 alkaline-responsive mRNAs including known bradyzoite genes (see Dataset S1) were changed by more than 1.5 fold in ^DD^AP2IV-3 tachyzoites grown in normal media plus 100 nM Shield 1. The log2 RMA values (x and y axis) for the 42 mRNAs were plotted: (A) CTG parent pH 7.0 VS pH 7.8, (B) CTG parent pH 7.0 VS CTG-^DD^AP2IV-3 pH 7.0, (C) CTG parent pH 7.0 VS CTG-^DD^AP2IV-3 pH 7.8. (D) Many mRNAs up regulated by the overexpression of ^DD^AP2IV-3 are repressed by ^DD^AP2IX-9 and no longer alkaline-stress inducible in parasites lacking AP2IV-3. Changes in mRNA levels of seven selected genes: in ^DD^AP2IV-3 CTG parasites grown at pH 7.0 in normal media plus Shield 1 (green bars and arrow), in ^DD^AP2IX-9 in parasites cultured in pH 8.2-media plus Shield 1 (red bars and arrow), and in Pru-*Δap2IV-3* parasites grown in pH 8.2 media without Shield 1 (blue bars and arrow). BAG1(TGME49_259020), LDH2(TGME49_291040), UNK(unknown gene TGME49_207210), PMA1(TGME49_252640), MIC UNK(TGME49_208740), DnaK(TGME49_202020).

### AP2IV-3 directly regulates transcription of the BAG1 gene

The global analysis of gene expression in AP2IV-3 transgenic strains demonstrates deletion or overexpression of AP2IV-3 strongly affects canonical BAG1 expression in a manner consistent with AP2IV-3 functioning as a transcriptional activator. To confirm AP2IV-3 directly regulates the BAG1 promoter, we generated a CTG transgenic strain co-expressing ^DD^AP2IV-3 and a second gene encoding firefly luciferase driven by the BAG1 promoter (BAG1-Luc). In previous studies, the BAG1 promoter fragment used here (1195bp, 5’-flanking the BAG1 coding region) conferred alkaline-stress inducible luciferase expression in developmentally competent strains (25). As a control, we also isolated a CTG transgenic strain expressing only the BAG1-Luc reporter. Results from luciferase assays using 100 nM Shield 1 (Fig. 5A) demonstrated that luciferase expression was higher in the CTG strain expressing ^DD^AP2IV-3 (yellow bars) at all pH conditions compared to the CTG parent strain only expressing BAG1-Luc (blue bars). This difference was strongly amplified by increasing alkaline-stress; CTG-^DD^AP2IV-3::BAG1-Luc parasites cultured in pH 8.2 media plus 100 nM Shield 1 expressed luciferase at ~20-fold higher levels than the CTG-BAG1-Luc control strain (Fig. 5A).The combination of increasing alkaline-stress plus Shield 1 also elevated native BAG1 protein levels in a Shield 1 dependent manner over the CTG parent control (Fig. 5A, inset Western Blot). These results demonstrated that ^DD^AP2IV-3 was not only regulating the BAG1-promoter driving the Luc cassette, but also the native BAG1 promoter.

**FIG 5.**
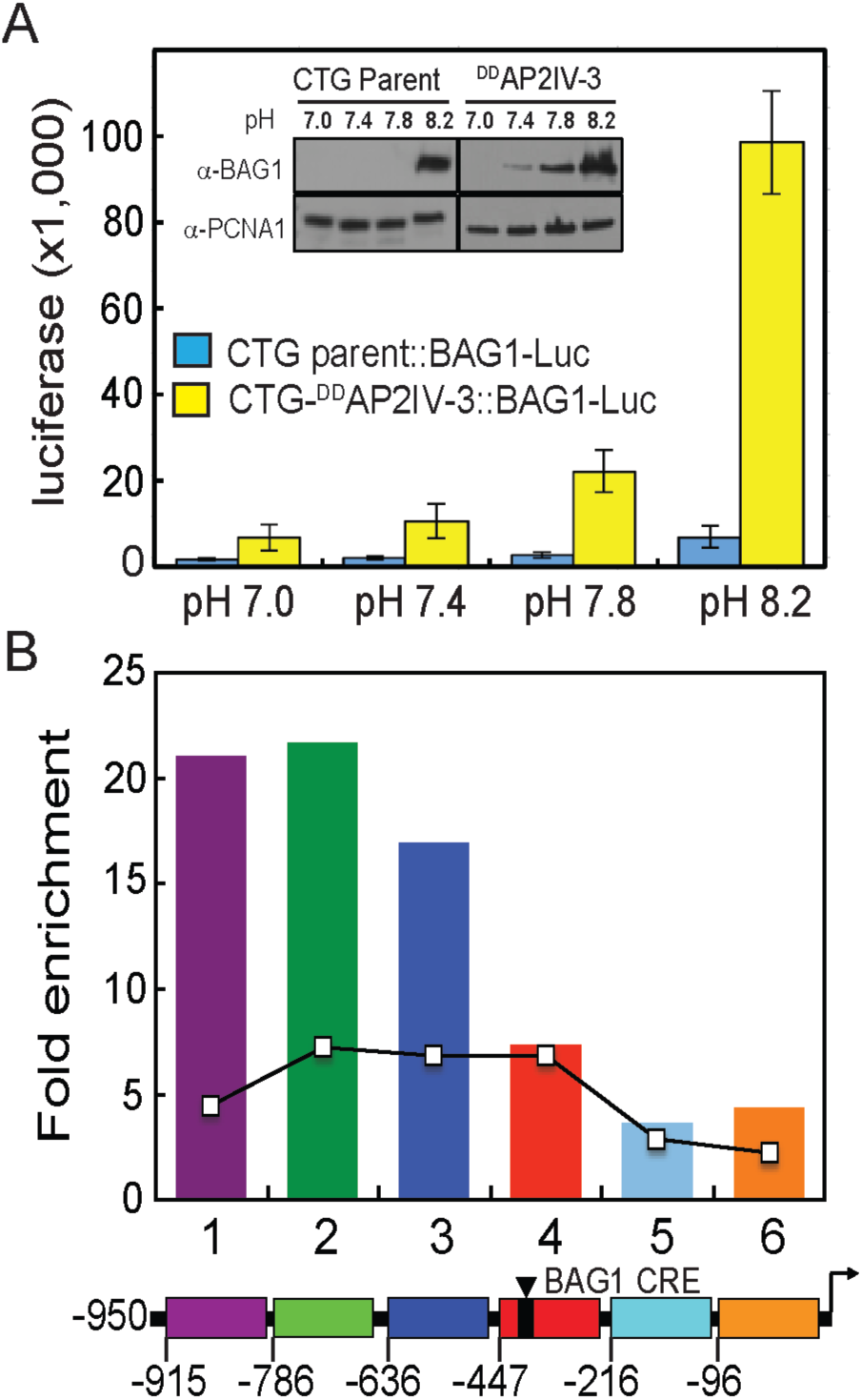
AP2IV-3 regulates BAG1 transcription. (A) Overexpression of AP2IV-3 induces the expression of the BAG1 promoter driving firefly luciferase (BAG1-Luc). CTG transgenic clones expressing ^DD^AP2IV-3 and BAG1-Luc (yellow bars) or BAG1-Luc alone (blue bars) were grown in media as indicated (pH 7.0-pH 8.2). Lysates from infected HFF cells were prepared and assayed for luciferase expression. Inset: Western analysis shows that Shield 1 overexpression of ^DD^AP2IV-3 induces the expression of native BAG1 protein as detected with anti-BAG1 monoclonal antibody. TgPCNA1 staining is included to demonstrate equal protein loading. (B) ^DD^AP2IV-3 occupies the native BAG1 promoter in parasite chromatin. Specific binding was monitored in specific sequences of the BAG1 promoter 5’ to the BAG1 coding region (-950 bp flanking including the BAG1 5’-UTR up to the coding ATG). The region of the BAG1 promoter between-950bp and-447bp (purple, green and blue regions) showed the highest enrichment and overlaps the regions where AP2IX-9 also binds as indicated by the overlaid line graph (6). Note that the BAG1 promoter element (BAG-CRE) previously mapped as required for Compound 1 induction (25) is located downstream of the regions of highest AP2IV-3 binding.

The direct regulation of the BAG1 promoter by AP2IV-3 was further validated utilizing chromatin immunoprecipitation followed by quantitative PCR (Fig. 5B). The binding of ^DD^AP2IV-3 to parasite genomic DNA demonstrated a specific preference for the BAG1 promoter. The pattern of ^DD^AP2IV-3 chromatin interaction overlaps the binding of AP2IX-9 to the BAG1 promoter (Fig. 5B, line graph)(6), although AP2IV-3 binding appeared to be stronger and to have a higher preference for the −950 to −636 region than AP2IX-9. Intriguingly, these regions of the BAG1 promoter are upstream of a key BAG1 promoter element previously mapped in Compound 1-treated Type II and III strain parasites (25).

## DISCUSSION

We have confirmed that two of the six *Toxoplasma* ApiAP2 factors (AP2IX-9 and AP2IV-3) induced early in bradyzoite development have important roles in regulating bradyzoite gene expression. AP2IX-9 is a transcriptional repressor that prevents the induction of bradyzoite gene expression and reduces tissue cyst formation (6), and thus, it was not surprising that disrupting AP2IX-9 in this study led to transgenic parasites that more readily formed tissue cysts under mild stress conditions. By contrast, AP2IV-3 is a transcriptional activator capable of enhancing tissue cyst formation when overexpressed or causing lower tissue cyst formation when deleted. Intriguingly, these two ApiAP2 factors regulate many of the same genes and chromatin immunoprecipitation and luciferase promoter assays clearly demonstrated AP2IX-9 and AP2IV-3 regulate the BAG1 promoter with opposite consequences and also specifically bind the BAG1 promoter in chromatin (Fig. 5B). We do not know if AP2IX-9 and AP2IV-3 directly compete or operate sequentially, but separately to control bradyzoite gene expression. In addition to AP2IX-9 and AP2IV-3, there is evidence for two other ApiAP2 factors regulating BAG1 gene expression (AP2XI-4, ref (27) and AP2IX-4, Huang and Sullivan, manuscript in preparation). Therefore, at a minimum, BAG1 transcription is controlled by four ApiAP2 factors; two activators (AP2XI-4, AP2IV-3) and two repressors (AP2IX-9, AP2IX-4). Our discovery of multiple ApiAP2 factors regulating BAG1 is consistent with our earlier investigation (25) that revealed the BAG1 promoter contains a number of potential repressor and activator promoter elements and multiple nuclear protein complexes could be detected binding BAG1 promoter DNA fragments. Further studies will be needed to precisely map the BAG1 promoter elements each factor binds and to determine whether these factors are contained in the same or different chromatin complexes.

Our group and others have established that bradyzoite development requires parasite replication (28), while also the slowing of the tachyzoite cell cycle primarily by increasing the length of the G1 phase (29). The shift to slower growth in spontaneous models of development is quite rapid occurring ~20 divisions following the emergence of the tachyzoite stage (29, 30). Importantly, early switching parasites have very similar transcriptomes to tachyzoites when compared to more mature bradyzoites (18) indicating these early developing parasites are largely slow growing tachyzoites, which can be considered pre-bradyzoite (Fig. 6). ApiAP2 factors likely have an important roles in regulating these growth to development transitions, and consistent with this idea, we have shown that the AP2IX-9 repressor promotes continued tachyzoite replication in Type II and III strains, while also preventing an increase in bradyzoite gene expression or production of tissue cysts (6). ApiAP2 factors may also influence the intravacuolar behavior of parasites. The cell cycle state and bradyzoite marker expression of individual parasites following alkaline-shift is asynchronous as compared to the tightly synchronous cell cycles of tachyzoites replicating in the same vacuole (21). The molecular mechanisms responsible for intravacuole tachyzoite growth synchrony and bradyzoite developmental asynchrony are not understood. However, it is clear that ApiAP2 factor expression follows this dichotomy and may be regulating these changes. Cyclical ApiAP2 factors in tachyzoites sharing the same vacuole are tightly coexpressed with specific ApiAP2 factor profiles providing markers for each cell cycle phase (12). By contrast, in alkaline-stressed differentiating populations, vacuolar ApiAP2 expression is no longer tightly coordinated. By dual tagging ApiAP2 factors in the same parasite, we have determined AP2IX-9 is generally up regulated first, while peak expression of AP2IV-3 coincides with the decline in AP2IX-9 expression (Fig. 1B). However, these expression patterns were population based not vacuole exclusive, as individual parasites within the same vacuole might express either or both nuclear factors. It is reasonable to suggest that it is likely to be more important for daughter tachyzoites to become infectious at the same time than for bradyzoites to become simultaneously dormant in the same tissue cyst, which may offer a biological rationale for the switch in ApiAP2 factor intravacuolar expression profile. There is also good biological logic to express an ApiAP2 bradyzoite repressor that promotes the tachyzoite stage earlier rather than later in the switching process as we observe for AP2IX-9 (6). Importantly, the differential expression of ApiAP2 factors during bradyzoite development now offers an opportunity to better understand the transitions in this developmental pathway at the level of the individual parasite.

**FIG 6.**
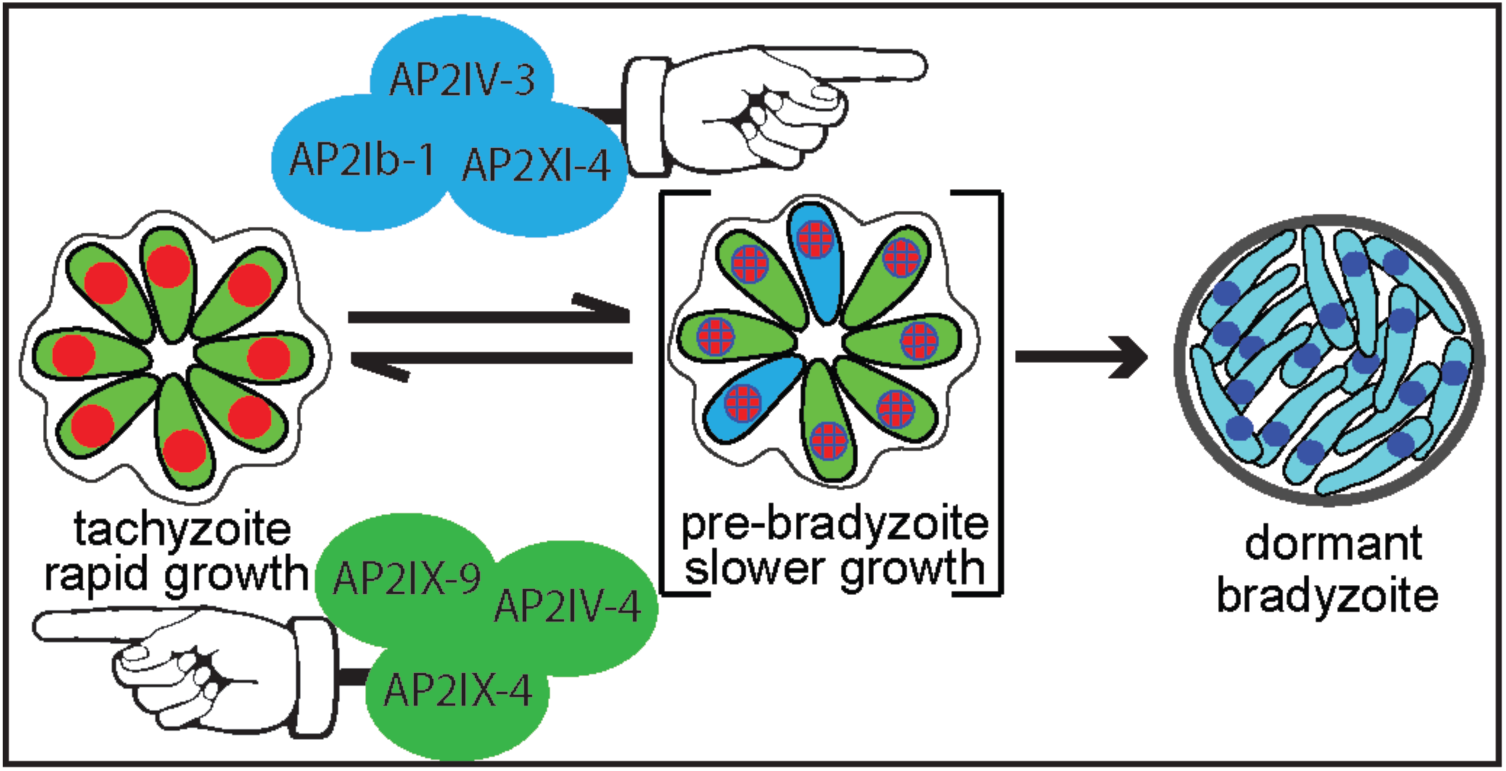
Model of ApiAP2 regulation of bradyzoite development.

A growing body of data indicate that fulfilling the biotic requirement of the intermediate life cycle is largely the province of the tachyzoite stage and not of parasites progressively engaged in developing and ultimately residing for the long term in the specialized tissue cyst. Oral infections by sporozoites or bradyzoites rapidly leads to population wide re-development of the tachyzoite stage, which is responsible for parasitema in animals (31, 32). A recent report demonstrates that the spread of tachyzoites into the vascular system in mice leads to tachyzoite infection of endothelial cells of the small capillaries of the brain that upon lysis travel across the blood brain barrier to provide the primary parasite source of brain tissue cysts (33). Consistent with this discovery, a recent large study of *Toxoplasma* development in mice demonstrates that tissue cyst size in brain tissue is related to the tachyzoite vacuole size at the time of switching (34) with average cyst size and number becoming stable after the first few weeks post-infection (34, 35). Increases in tissue cyst size in later stages of chronic infection is largely independent of parasite replication (34) and there is little evidence that cycles of cyst rupture amplify tissue cyst numbers, as these events are rare and minimally productive (0.26% of brain cysts recrudesce)(36). Some replication of bradyzoites or pre-bradyzoites can be detected well into development (34) and there is evidence tachyzoites formed from sporozoites are capable of switching to bradyzoites after very few divisions if artificially stressed (30). However, the contribution of mature bradyzoite replication to the biotic expansion of *Toxoplasma* in infected animals is likely very modest. Our recent discoveries of multiple ApiAP2 repressors and activators regulating growth and tissue cyst formation provide substantial support for the close relationship between the tachyzoite cell cycle and mechanisms that trigger the progressive development of the tissue cyst. In addition to the AP2IX-9 and AP2IV-3 factors studied here, our group recently identified two cell cycle regulated ApiAP2 factors of the tachyzoite whose major functions are to repress bradyzoite gene expression (Radke, Huang, Sullivan, and White, manuscripts in preparation). Thus, there is now direct evidence for at least six ApiAP2 factors regulating bradyzoite development that act at the interface of tachyzoite growth and the stress-responses that trigger the bradyzoite developmental pathway (Fig. 6 model). These results indicate that the competition between continued growth against the development of dormant tissue cysts required for transmission likely drove the evolution of developmental transcriptional mechanisms in *Toxoplasma*. Further, this model is consistent with our previous work indicating bradyzoite differentiation is likely the default pathway of life cycle competent strains (29, 30), and therefore, one role of ApiAP2 bradyzoite repressors is to ensure reproduction of tachyzoites meets minimum biotic requirements. Opposing transcriptional forces is a common adaptive feature of gene expression mechanisms from bacteria to man (37), and is a particular theme of AP2 regulation of plant gene expression (38). Plant AP2 repressors serve specific roles in flower development (39) and the response to environmental stress (40) and there appears to be two kinds of AP2 repressors in plants; an active form that binds DNA directly and a second type of repressor that operates by inhibiting activator activity (38). Our studies of ApiAP2 factors may have uncovered both types of repressors in *Toxoplasma* (6)(Huang and Sullivan, manuscript in preparation)

It is early in our understanding of how ApiAP2 factors function in *Toxoplasma*, and other Apicomplexans, although there is already more complexity in coccidian ApiAP2 mechanisms than what might have been expected. Coccidian parasites have more ApiAP2 factors than other Apicomplexan parasite families, such as the *Plasmodium spp*, that can partially be explained by a larger genome and transcriptome. Another possible explanation is the wider intermediate host ranges of coccidian parasites, and this may explain the relatively few ApiAP2 factors exclusively expressed in the feline definitive life cycle (Table 1). In other eukaryotes including mammals, cell division is a requirement for subsequent development as the process of chromosome replication allows for a specialized transcriptional programming of terminally differentiated cells (41, 42). Intriguingly, *Toxoplasma* appears able to tap into these mechanisms to find long-lived host tissue to develop the tissue cyst (43). It will be interesting to determine whether host-parasite signal transduction mechanisms acting through ApiAP2 factor mechanisms are responsible for achieving the coordination of tissue cyst formation in terminally differentiated host cells. There are clearly candidates for host factors involved in this communication pathway that negatively regulate host cell growth such as CDA-1, p21, and cyclin B1 (1, 43, 44) and *Toxoplasma* kinases that are likely involved such as casein kinase (1) and TgPKA (45). Therefore, ApiAP2 factors are intriguing candidates that could receive critical signals from these and other host pathways and drive the appropriate developmental transcriptional response within the parasite.

## MATERIALS AND METHODS

### Cell culture and transgenic strains

Parasite cultures were maintained in primary human foreskin fibroblasts (HFF) as previously described (46). All genetically modified strains (see full list of parent and transgenic strains in Dataset S3) generated here were grown in confluent monolayers of HFF cells in medium composed of Dulbecco’s modified Eagle Medium (DMEM) with 4.5 g/L Glucose, L-glutamine & Sodium pyruvate (Corning Cellgro), 5% fetal bovine serum (FBS) (Sigma) and 1% Pen/Strep/Amphotericine B (Hyclone) at 37°C with 5% CO_2_.

### Overexpression of AP2IV-3

For conditional overexpression, the AP2IV-3 single exon coding region was PCR amplified from genomic DNA to incorporate in frame Xma1/Sbf1 sites, which were used to clone the purified PCR fragment into the pCTDDHA3x plasmid. This cloning results in the fusion of the FKBP peptide (11.2 kDa) and 3 copies of the HA epitope for detection (4.4 kDa) in frame with the N-terminus of AP2IV-3 protein with a final fusion protein mass of ~162 kDa (designated ^DD^AP2IV-3). The plasmid pCTDDHA3x-AP2IV-3 was introduced by electroporation into the Type III CTG strain. Transgenic parasites were selected using chloramphenicol (20 μM) and clones were isolated by limiting dilution. Overexpression of AP2IV-3 was achieved by adding 100 nM Shield 1 synthesized in house to culture media.

### AP2IV-3 and AP2IX-9 knockout and complementation

For disruption of ApiAP2 factors, we have used the multi-gRNA CRISPR/Cas9 system. The gRNAs for each gene was generated by CRISPR gRNA design tool supported from DNA2.0 (www.dna20.com). The CRISPR cas9-gRNA plasmid (pSAG1::Cas9-U6::sgUPRT) was provided by David Sibley (Washington University, St Louis). The single gRNA in this plasmid was replaced by a short oligonucleotide matching each targeted gene (see Dataset S3 in the supplemental data) using Q5 DNA site directed mutagenesis (NEB, Ipswich, MA) with a common reverse primer (5’AACTTGACATCCCCATTTAC3’). Gene specific ApiAP2 CRISPR-Cas9-gRNA plasmids (Dataset S3) were sequenced to confirm the correct sequence of the gRNA. To generate clean knockouts in *Toxoplasma* tachyzoites, two gRNA plasmids for AP2IV-3 and four gRNA plasmids for AP2IX-9 were designed in the same gene. A plasmid named 3Frag-AP2KO/DHFR-TS containing a pyrimethamine resistance cassette was constructed using 3-fragment gateway (Thermal Fischer Scientific, Waltham, MA). Both the 5’ and 3’ UTR of ApiAP2 genomic coding regions were PCR amplified from Type I genomic DNA and used to create Gateway entry clones. The 5’UTR was cloned into the pDONR_P4-P1r vector and the 3’UTR was cloned into the pDONR_P2r-P3 vector by the BP recombination reaction. After isolation of the entry clones, pDONR_P4-P1r_AP2 and pDONR_P2r-P3_AP2 were combined in a LR recombination reaction with p221-DHFR and pDEST_R4-R3 vectors to create pDEST_AP2_DHFR knockout plasmid. The final plasmids were introduced into Pru and CTG strains and transgenic parasites were selected using pyrimethamine followed by clonal isolation by limiting dilution. The 30 μg of each gRNA plasmids and pDEST_AP2_DHFR knockout plasmid were transfected into 3 x 10^7^ parasites by using electroporation. For complementation of AP2IV-3, purified cosmid (PSBLJ40) carrying a single copy of the AP2IV-3 gene was transfected into Pru-*Δap2IV-3* parasites. To quantify genetic rescue, established drug-resistant populations were tested for tissue cyst formation in alkaline media (pH 8.2) and AP2IV-3 mRNA as well as control GAPDH mRNA were analyzed by quantitative RT-PCR.

### Immunofluorescence assay and Western analysis

Parasites were inoculated on confluent HFF coverslips for the indicated times and IFA assays performed by using the following primary reagents: anti-HA antibody (Roche; rat mAb 3F10;1:500), anti-myc antibody (Santa Cruz Biotechnology, mouse mAb, 1:500), biotin-labeled Dolichos biflorus agglutinin (1:3000, Vector labs, CA), and DAPI (0.5mg/ml). All Alexa (Molecular Probes, CA) and streptavidin (Vector Labs, CA) conjugated secondary antibodies were used at 1:1000. Image acquisition was performed on a Zeiss Axiovert microscope equipped with 100x objective. Western blotting with specific antibody monitored the protein expression in *Toxoplasma.* Purified parasites were lysed in SDS-PAGE sample buffer with Leammli loading dye, heated at 95°C for 10 min, and briefly sonicated. After separation on the gel, proteins were transferred onto nitrocellulose membrane and detected with monoclonal antibodies and horseradish peroxidase (HRP) conjugated secondary antibodies (Jackson ImmunoResearch, PA) and visualized using enhanced chemiluminescence detection (PerkinElmer).

### Microarray analysis

Total RNA was purified from intracellular CTG, CTG-^DD^AP2IV-3, and CTG-*Δap2IV-3* parasites maintained at pH 7.0 and pH 7.8 media (32-36 hours post-infection) or following 48 hours exposure to pH 8.2 media using RNeasy Mini kit (Qiagen) according to manufacturer’s instructions. Two biological replicates were prepared for each parasite line under each experimental condition and RNA quality was assessed using the Agilent Bioanalyzer 2100 (Agilent Technology).Following fragmentation, 10 μg of cRNA was hybridized for 16 hours at 45 °C to the *Toxoplasma* affymetrix U133 Plus 2.0 arrays. GeneChips were washed and stained in the Affymetrix Fluidics Station 450. Hybridization data was analyzed using GeneSpring GX software package (v12.6.1, Agilent) and all data were deposited at NCBI GEO (GSE89469).

### Luciferase assays

Luciferase assays were performed according to manufacturers protocols (Promega, Madison, WI) with modifications as previously described (25). Briefly, HFF cells in T25cm^2^ flasks were inoculated with CTG-BAG1-Luc and CTG-^DD^AP2IV-3::BAG1-Luc transgenic parasites at 3:1 MOI, parasites were allowed to invade for 1 h, and the culture media changed to either pH 8.2 or pH 7.0 media plus or minus 100nM Shield 1. The alkaline-shifted cultures were grown in non-CO_2_ conditions for 48 h and then harvested for whole cell lysates according to manufacturers protocols. Each experimental condition was assessed in three independent cultures.

### Chromatin Immunoprecipitation and qPCR

Parasites were inoculated at 3:1 MOI in T175 cm^2^ flasks, allowed to invade for 1h, rinsed three times with HBSS to remove free floating parasites, and media was replaced with pH 8.2 media supplemented with 100 nM Shield 1. Chip-qPCR was performed by published methods (6).

## Acknowledgements

We are grateful for the discussions and contributions to these studies from William Sullivan Jr, Kami Kim, Elena Suvorova, and Louis Weiss. We thank David Sibley for kindly sharing CRISPR reagents and we are in debt to the genomics core facility of the Moffitt Cancer Center who performed microarray hybridizations and data extraction for these studies. Finally, we want to thank the folks at ToxoDB that have generated and maintain one of the highest quality molecular repositories for eukaryotic pathogens. This work was supported by grants from the National Institutes of Health to MWW (R01-AI077662, R01-AI089885, and R56-AI124682).

